# Expression and Splicing Mediate Distinct Biological Signals

**DOI:** 10.1101/2022.08.29.505720

**Authors:** Søren Helweg Dam, Lars Rønn Olsen, Kristoffer Vitting-Seerup

**Author notes:** **Contact** SHD LRO KVS.

## Abstract

**Background:** Through alternative splicing, most human genes produce multiple isoforms in a cell-, tissue-, and disease-specific manner. Numerous studies show that alternative splicing is essential for development, diseases and their treatments. Despite these important examples, the extent and biological relevance of splicing are currently unknown.

**Results:** To solve this problem, we developed pairedGSEA and used it to profile transcriptional changes in 100 representative RNA-seq datasets. Our systematic analysis demonstrates that changes in splicing, on average, contribute to 48.1% of the biological signal in expression analyses. Gene-set enrichment analysis furthermore indicates that expression and splicing both convey shared and distinct biological signals.

**Conclusion:** These findings establish alternative splicing as a major regulator of the human condition and suggest that most contemporary RNA-seq studies likely miss out on critical biological insights. We anticipate our results will contribute to the transition from a gene-centric to an isoform-centric research paradigm.

## Background

High-throughput sequencing of RNA (RNA-seq) has revolutionized our understanding of molecular biology. It has become the standard approach for unraveling the complexity of living organisms, as illustrated by tens of thousands of already published RNA-seq datasets[1, 2]. One of the primary uses of RNA-seq is to compare samples in case/control settings by analyzing differential gene expression and differential gene splicing. Regardless of whether differential expression or differential splicing is analyzed, the result is that hundreds to thousands of genes are found to change significantly between conditions. Most scientists use gene-set enrichment analysis (GSEA) to interpret these long gene lists. Hence GSEA has become a cornerstone within genomics because of its ability to extract biologically meaningful insights.

Differential splicing is caused by changes in how a pre-mRNA is spliced into a mature mRNA molecule. RNA splicing is an essential molecular mechanism that removes introns from precursor mRNA, joining exons to form the mature mRNA. Along with alternative transcription start and termination sites, alternative splicing (hereon jointly referred to as splicing) enables genes to give rise to multiple transcripts. During the past decades, this has changed how we think of genes. A gene is no longer viewed as encoding just one transcribable product but instead as giving rise to several mRNA transcripts. It is estimated that 92–94% of human genes undergo splicing [3, 4] and that protein-coding genes produce, on average, seven transcripts[5]. However, recent analyses of long-read data suggest that this is an understatement[6, 7]. In addition, recent large-scale studies find that the splicing of most human protein-coding genes is affected by naturally occurring genetic variation (SNPs)[8–10] suggesting further transcript diversity.

Splicing often induces significant functional alterations of the gene product. For one, splicing is essential for defining cell types[11], e.g., the difference between naïve and activated memory T cells, which are defined by the two CD45 isoforms CD45RA and CD45RO, respectively[12]. The switch to the CD45RO isoform drastically increases the sensitivity of the T cell receptor [13]. Besides being essential for defining cell types, splicing is also central to cellular functions, such as apoptosis. Many apoptotic proteins, such as BLC-X[14] and caspase-2[15], exist in two opposing isoforms, one pro-apoptotic and one anti-apoptotic. As part of the apoptotic cascade, these genes switch from producing the anti-apoptotic to the pro-apoptotic isoform, thereby simultaneously removing the breaks and accelerating apoptosis[15].

Although most scientists analyze their RNA-seq data using differential gene expression followed by GSEA, splicing is generally overlooked. Our literature analysis indicated that only 12% of articles that analyzed RNA-sequencing data in 2020 did any sub-gene level analysis (Figure S1). However, an exciting pattern emerged from recent studies jointly investigating differential expression and splicing: The overlap between the differentially expressed and differentially spliced genes is low [6, 10, 16–20]. This suggests that expression and splicing could mediate distinct biological signals. However, such studies are merely individual examples; to our knowledge, no systematic analysis of splicing currently exists. Thus, we do not know how widespread or biologically relevant changes in splicing are.

Here, we systematically analyze and compare differential expression and differential splicing in human genes. We chose to focus on bulk RNA-seq because it is currently the only widely available high-throughput approach where both the technology and analysis approaches are mature enough to analyze splicing. Across 100 representative RNA-seq datasets, we see that changes in splicing are pervasive and, compared to differential expression, mediate both shared and distinct biological signals. Our findings indicate that many scientists are underutilizing their RNA-seq data, thereby missing important biological insights.

## Results

### pairedGSEA enables paired analysis of differential expression and splicing

We assembled a robust analysis pipeline to compare differential splicing and expression systematically. The analysis pipeline is implemented in the R package pairedGSEA (Figure 1A) (https://bioconductor.org/packages/pairedGSEA) and enables easy:

**Figure 1:**
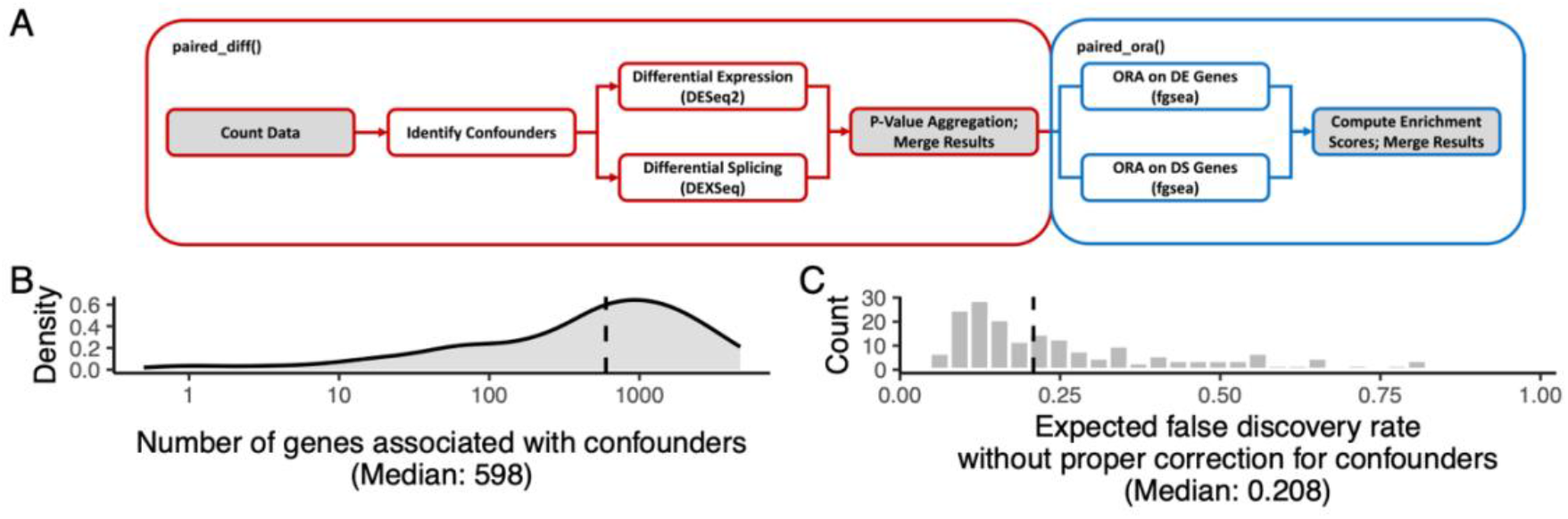
pairedGSEA and confounder-mediated false discoveries. **A**) Flowchart of the pairedGSEA R package and its functions (red and blue rounded squares). The gray and white backgrounds in the boxes indicate data and functionality, respectively. **B**) The distribution of false positives, i.e., the number of significantly differentially expressed genes only found when not corrected for confounders, across the 199 comparisons. **C**) Histogram of the false discovery rate when not correcting for confounders. Significance is defined as having an FDR-adjusted p-value of <0.05.

- [21]Identification of unknown unwanted experimental (confounding) factors, such as batch effects (via sva [21]). These, along with user-supplied covariates, are propagated into the differential analysis thereby ensuring they do not affect the downstream analysis.
- Analysis of differential splicing and differential expression (via DEXSeq [22] and DESeq2 [23], respectively)
- Analysis and comparison of the result of both expression and splicing for over-represented gene sets (via fgsea [24]).

To utilize the pipeline, we curated 100 randomly selected high-quality RNA-seq datasets (Table S1, see Methods) covering a wide range of study types. Examples include using different growth media, inhibition or overexpression experiments, and analyzing the effects of treating various human diseases with different drugs (Supplementary Table S1). Importantly the random selection ensures our results represent what scientists can expect from their own data when performing a new experiment as it reflects cell, tissue, and treatment preferences in published data. We applied pairedGSEA to all 100 datasets resulting in 199 comparable analyses of differential expression and differential splicing, as some datasets had more than two conditions (Supplementary data). Interestingly, we found that every dataset had genes significantly affected by confounding factors (median 598 genes, Figure 1B). Indeed, if these confounders were not considered, the false discovery rate (FDR) would be ∽20.8% on average instead of the expected 5% (Figure 1C). In other words, confounders like batch effects are pervasive, and if not considered, the results of differential analysis will essentially be unusable.

### Differential Splicing is just as Frequent as Differential Expression

To systematically compare differential splicing and differential expression, we extracted the significant results (FDR Corrected P-values < 0.05) from all 199 comparisons. Across all comparisons, we, on average, found 4327 (26.1% of tested) genes that were significantly differentially expressed, and 2247 (12.9% of tested) genes were significantly differentially spliced (Figure 2A-B). Among these, on average, 1252 genes were significant in both analyses. This means that, on average, 33.6% of the differentially expressed genes were also significantly affected by splicing (Figure 2C). Notably, the splicing changes were not just due to lowly expressed isoforms; the changing isoforms contributed, on average, 45.6% of the parent gene expression (Figure 2D). Thus, more than one in three significantly differentially expressed genes contained splicing differences that could dramatically change the gene function, e.g., through dominant negative splice variants, highlighting the need to consider splicing changes.

**Figure 2:**
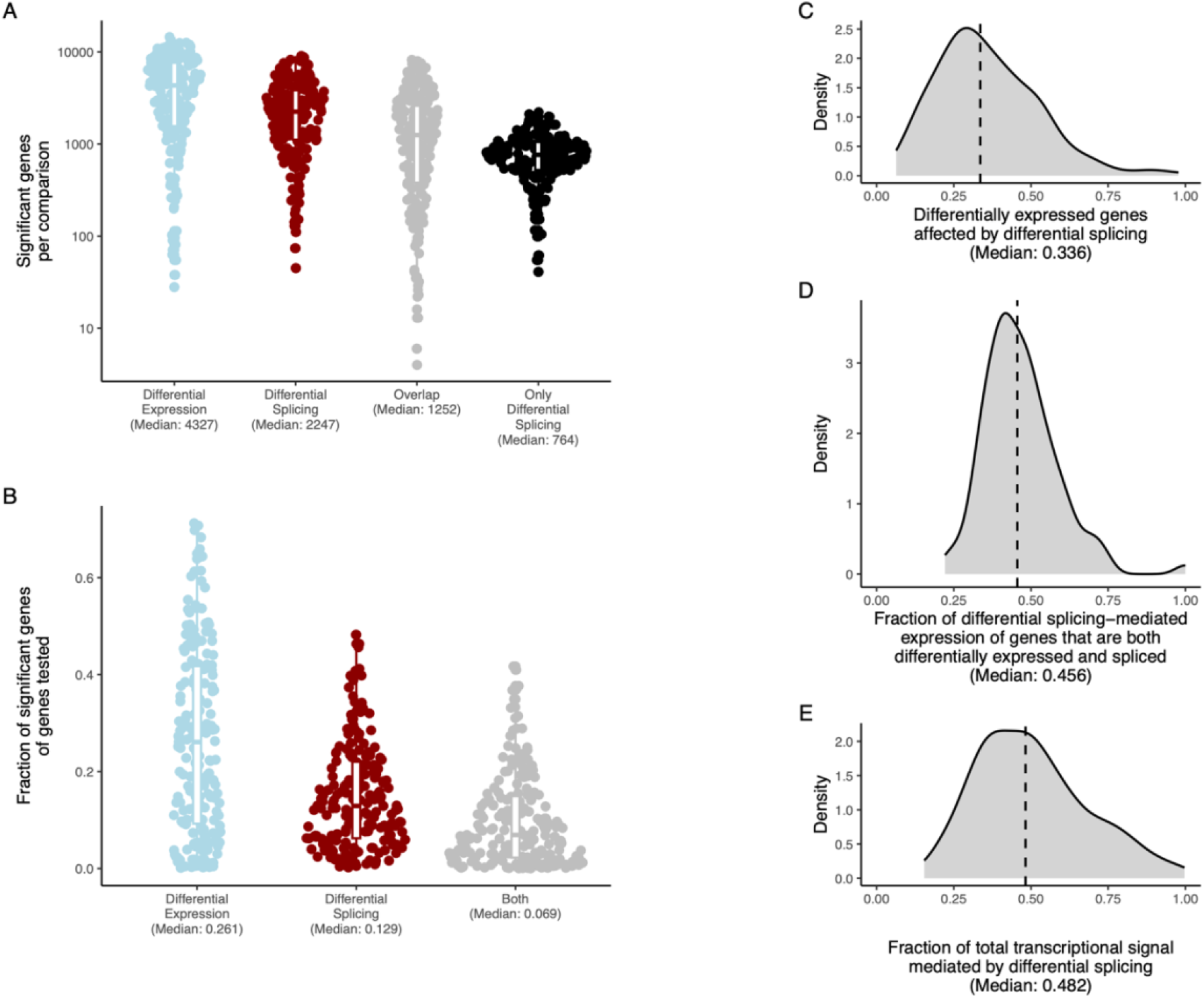
Differential Splicing is just as Frequent as Differential Expression. **A**) The number of significant genes for each comparison across analyses. **B**) For each analysis, the fraction of genes tested that were deemed significant. **C**) The fraction of differentially expressed genes that were also differentially spliced. **D)** Within the genes that are both differentially expressed and spliced, we calculated the fraction of the gene expression that is contributed by differentially spliced transcripts. For each analysis, we extracted the median. **E**) The number of differentially spliced genes as a fraction of the total number of genes either differentially spliced or expressed genes (total transcriptional signal). Across all panes, significance is defined as having an FDR-adjusted P-value of <0.05. Medians are indicated for all plots.

Across the 199 comparisons, our systematic analysis found that 48.2% of the combined biological signal was at least partially mediated by changes in splicing (Figure 2E). Thus, it is not surprising that 93.0% of all multi-isoform genes were significantly differentially spliced in at least one dataset, indicating that most genes utilize splicing as part of adapting to new circumstances.

In summary, this indicates that splicing is an integral part of the response pattern for all genes on par with expression changes.

### Splicing and Expression Regulate Distinct Biological Processes

Since genes are rarely independent functional entities, we used gene-set enrichment analysis to analyze gene sets enriched among differentially spliced or expressed genes. This reduced the result of the differential expression significant to, on average, 1829 significant gene sets and the differential splicing result to, on average, 32 gene sets (Figure 3A). On average, 21 gene sets overlapped between analyses, indicating that many cellular circuits are regulated through both expression and splicing. The remaining gene sets were solely modulated through differential splicing, again suggesting that splicing can mediates distinct biological information (Figure 3A).

**Figure 3:**
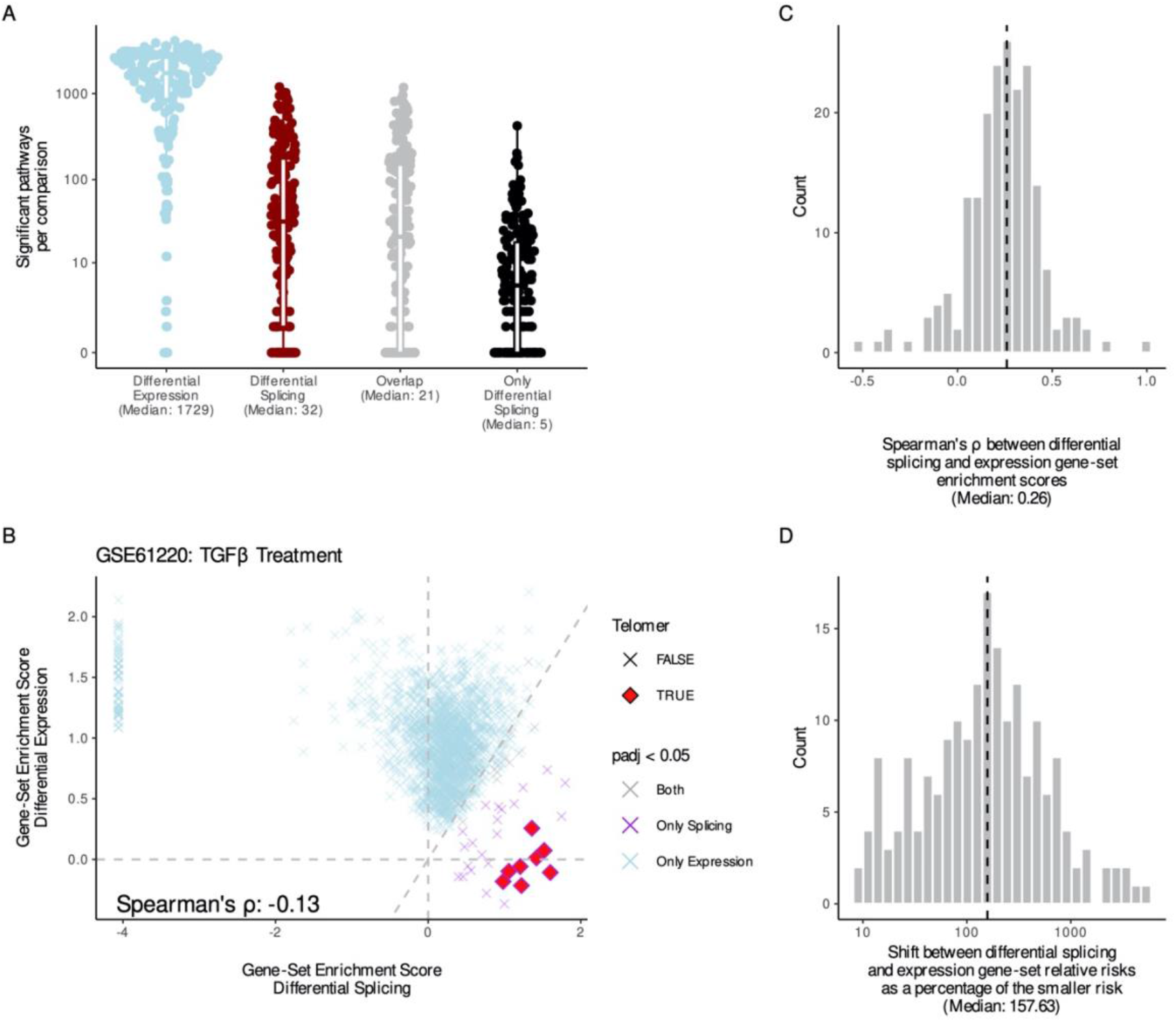
Splicing and expression regulate distinct biological processes. **A**) The number of gene sets significantly enriched among genes from either analysis across comparisons. **B**) Results from the Tian *et al*. [25] study showing the gene-set enrichment scores of gene sets enriched among the differentially spliced (x-axis) and differentially expressed (y-axis) genes. Only gene sets significantly enriched among differentially spliced or differentially expressed genes (indicated by color) are shown. The shape highlights gene sets where the name contains the word “telomer”. Spearman’s correlation is indicated in the lower left corner. **C**) Histogram of the Spearman’s correlations between gene-set enrichment scores for gene sets significantly enriched among differentially expressed or spliced genes. **D**) For each comparison, we calculated the median differences between the relative risks of gene sets enriched among differentially expressed and spliced genes as the percent change of the smallest risk score.

To illustrate this point, we analyzed the study by Tian *et al*. [25], where human small airway epithelial is treated with TGFβ: a known regulator of telomere length [26, 27]. Surprisingly, the gene-set enrichment analysis showed no significant association between differentially expressed genes and telomere-related gene sets. However, the telomere-related gene sets were significantly enriched among the differentially spliced genes (Figure 3B), indicating that the regulation of telomeres by TGFβ is solely mediated through changes in splicing [28].

If a gene set is regulated by both expression and splicing (shared regulation), the enrichment score for the splicing and expression-based enrichment analysis is expected to be similar (supplementary figure 4). Intriguingly, we find that the gene-set enrichment scores for the splicing and expression gene-set enrichment analysis had a slightly negative correlation in the TGFβ study (Spearman’s ρ of -0.13, Figure 3B). Inspired by this, we analyzed the correlation of gene-set enrichment scores for all significantly enriched gene sets within each of the 199 comparisons. We found that gene-set enrichment scores of splicing and expression changes were mostly uncorrelated (Median correlation: 0.26, Figure 3C). In addition, the relative risks were also substantially different (Figure 3D), suggesting substantial regulation differences. These observations also held true when considering gene sets enriched among either differentially expressed or differentially spliced genes (Figure S2-3). Interestingly, the gene sets significantly enriched with both differentially expressed and spliced genes seemed to act differently. They had much smaller enrichment score differences and a Spearman’s correlation of 0.68 (Figure S2-3). This similarity suggests co-regulation of these pathways. Thus, our systematic analysis across 100 representative RNA-seq datasets indicates that differential expression and differential splicing act both independently and jointly to regulate biological function.

## Discussion

In this study, we have used 100 representative RNA-seq datasets to show that differential splicing can significantly affect almost all multi-isoform genes. Nearly half of the observed biological signal originates at least partly from splicing, indicating that changes in splicing are an integral part of biological responses. Since we estimated that 88% of recent articles about RNA-seq are not doing any sub-gene-level analysis, our results indicate a considerable opportunity gap. Reanalyzing these studies might reveal many missed discoveries important for both the fundamental and the translational sciences.

Our results are obtained through a systematic analysis of RNA-seq data. Therefore, some observed changes could exist without being translated to the protein level. Many research groups have tried to determine how much splicing is reflected at the protein level, e.g., using bottom-up proteomics, resulting in a lively debate on the topic. Unfortunately, resolving this mystery is hindered by several factors, such as high sequence similarity of protein isoforms and low coverage of proteomics [29].

Furthermore, peptides generated by standard bottom-up proteomics are heavily depleted for peptides that cross exon-exon junctions, making it harder to detect splicing differences [30]. The result of the discussion is that there are multiple papers on either side of the argument, respectively, concluding that there is a lack of proteomics evidence for splicing [31, 32] or ample proteomics evidence for splicing [33–35]. This disagreement is highlighted by commentary papers challenging articles on both sides of the argument [36, 37]. The central point in this discussion seems to be how to produce trustworthy but not overly conservative results; unfortunately, no consensus seems to be on the horizon.

Until such consensus is reached, we believe it is worth looking at other data types to estimate the abundance of splicing in proteins. For instance, top-down proteomics does not suffer from the above-described shortcomings and easily identifies and distinguishes between protein isoforms [38, 39]. Prominent examples are Tran *et al*. [40] and Yu *et al*. [41] who identified isoforms for 22.4% and 46.5% of the proteins they identified (1043 and 628, respectively; our calculations, see Methods). Using RNA-seq, it has previously been determined that lab-validated changes in isoform expression are 16 times more frequent than expected by chance [42], indicating that transcriptional approaches do indeed capture protein-level changes. In addition, a growing number of studies are specifically designed to detect and analyze the impact of changes in protein isoforms. Large-scale studies of protein isoforms using technologies such as yeast two-hybrid have shown that the interaction partners for isoforms are very different from each other [43, 44]. Furthermore, several research groups find good correspondence between transcriptional and protein changes when analyzing isoform-targeted proteomics and Ribo-seq [39, 45–47]. Given this evidence, we and others find it highly likely that the majority of the transcriptional variation we described here will also be reflected in the proteome[48].

The analysis presented in this paper is based on the state-of-the-art bioinformatics tools DESeq2 [23] and DEXseq [22]. Reassuringly all our conclusions were replicated when we applied another widely used statistical method, voom-limma[49, 50], for identifying differentially expressed and spliced genes (Supplementary Figure 5,6,7). Despite the high performance of DEXSeq/limma it is well established that differential expression has more statistical power compared to differential splicing [51]. This means, that our estimates of the frequency and importance of splicing are probably underestimated.

Most systems biology relies on databases of annotation to make biological inferences. Almost all such databases, especially the most frequently used ones, only contain gene-level annotation. Since we find splicing is both frequent and significant, our tool provides an essential way to extend the functionality of these databases. Interestingly some people have started to work on transcript-level databases [52– 54], but there is still much to do. Another peculiarity that needs more investigation is the apparent inconsistency between the number of differential spliced genes and differentially spliced gene sets. This finding emphasizes the necessity for additional examination and exploration of the intricate relationship between expression and splicing regulation.

Our results show that splicing changes impact all genes with biologically relevant effects. Thus, it is clear that splicing should be considered when possible. It also highlights a massive caveat with technologies relying on only capturing the 5’ or 3’ end of transcripts. This is especially true for single-cell RNA-seq, where most datasets cannot be used to assess changes in splicing. In further support, the few single-cell datasets that can be used to analyze splicing show that splicing analysis leads to novel findings, including novel cell types [11].

## Conclusion

Our results indicate that the biological role of splicing is *on par* with the importance of changes in gene expression, similar to what has previously been estimated from the analysis of genetic data[10]. This has significant implications for most aspects of life sciences and equally affects wet labs doing mechanistic single-gene work and consortia investigating population-level genomics. Ultimately, utilizing the mostly untapped information hidden in differential splicing could pave the way for new clinical strategies within disease diagnosis, therapy, and precision medicine. For instance, detecting a switch in isoform usage in known oncogenes could indicate cancerous activity, cancer-specific isoforms could pose as targets for novel immunotherapies [55], and expression of specific isoforms could increase the risk of relapse or even render some therapies ineffective [56]. All sciences would benefit from updating how they work, moving from the current gene-centric research paradigm toward a more modern isoform-centric one.

## Methods

### Systematic literature survey

For the systematic literature survey, we used the same approach as in Vitting-Seerup *et al*. [42], except this time, we analyzed articles from the first six months of 2020. Briefly, we used PubMed to search for articles about RNA-seq and isoforms. We randomly selected 50 papers only found in the RNA-seq search and 50 articles also found in the isoform search. Then, we manually profiled how the data was analyzed, including whether transcript quantification was done and whether any sub-gene level analysis was done. The results were extrapolated to the expected fractions of all articles about RNA-seq data as described in Vitting-Seerup *et al*. [42].

### Curating representative datasets

To construct a collection of high-quality datasets, we considered >14,500 human RNA-seq studies [1]. From these, we computationally extracted high-quality datasets containing at least two conditions. High-quality datasets were defined as follows: 1) All samples within a dataset were predicted as bulk RNA-seq data with >90% certainty. 2) All samples having at least 40% of reads aligned to the human transcriptome. 3) All samples have at least 10 million aligned reads. 4) The study has a maximum of 50 samples (as DEXSeq does not scale well with increasing sample size). Next, we randomly selected 100 RNA-seq datasets and manually determined which groups to compare, extracting maximum four comparisons per dataset. The result was 199 comparisons from 100 datasets (Table S1).

### The pairedGSEA R package

A comprehensive guide on the functionalities of pairedGSEA is available in the package vignette (https://bioconductor.org/packages/pairedGSEA). The purpose of pairedGSEA is to make a baseline vs. case paired differential expression and splicing analysis simple. It assumes you have already preprocessed and aligned your sequencing data to obtain transcript level counts. Running pairedGSEA with default settings will filter out lowly expressed transcripts and detect potential batch effects (or other confounders) using sva [21]. If co-founders are detected, they will be added to the design matrix used for the differential analyses. pairedGSEA will then compute the differential expression and differential splicing on transcript level using DESeq2 [23][22] and DEXSeq [22][23], respectively. DESeq2 is run using a likelihood-ratio test using a reduced model where the condition information is removed. The results are extracted with a baseline vs. case contrast. The model used in DEXSeq adds the interaction between the transcript counts and the condition, and the confounding variables. DEXSeq does not allow a definition of a baseline as that is more abstract in differential splicing; however, pairedGSEA ensures the log fold changes correlate between the two analyses.

<u>As an alternative to DESeq2/DEXSeq, pairedGSEA can also do the differential analyses using limma. Here, a linear model is fitted using the same design matrix as for DESeq2. For differential expression, empirical bayes statistics are computed, while for differential splicing the fit is tested for log-fold-changes between transcripts of the same gene using the diffSplice function.</u>

In the final step of the paired differential expression and splicing analysis, pairedGSEA aggregates the transcript p-values to gene level using Lancaster aggregation [57] with base means as weights. The differential expression log fold changes are aggregated using a weighted mean with base means as weights. Again, the log fold change of differential splicing is a bit more abstract; therefore, it was chosen to keep the log fold change of the transcript with the lowest p-value as the log fold change of the corresponding gene. Then, the p-values are adjusted separately for the two analyses by false discovery rate using the Benjamini-Hochberg procedure, and the two results are merged into a single object. The results can then be directly used in the GSEA part of pairedGSEA. Significant genes are extracted by a user-defined adjusted P-value cutoff (defaulted to 0.05) and two over-representation analyses (ORA) are run using the fora function from the fgsea package [24]: one for differentially expressed genes and one for differentially spliced genes. Each ORA analysis is done with a separate background (universe) reflecting that single-isoform genes cannot be tested for differential splicing. Genes found in both will be included in both over-representation analyses. But before doing so, a list of gene sets is needed. pairedGSEA provides a function to extract gene sets from MSigDB [58], but users may use any database they prefer. After running the over-representation analyses, pairedGSEA computes an enrichment score for each gene set as the log2 relative risk. Specifically, it is calculated as: Log2((overlap / gene_set_size) / (significant_genes / total_genes_analyzed) + 0.06).

### Analysis of curated dataset

All data sets were obtained from a local copy of the ARCHS4 v11 database of transcript counts [1]. The Genome Reference Consortium Human Build 38 was used to retrieve transcript-to-gene associations, minimizing the number of lacking associations otherwise found in the ARCHS4 database. The baseline and case condition, and experiment titles were retrieved from the manually curated metadata described in the Data selection section. Paired differential analysis was run with pairedGSEA using default settings. The inbuilt wrapper for MSigDB extraction was used to create a list of gene sets from the ‘C5’-collection of Homo Sapiens gene sets, which were subsequently used in the pairedGSEA over-representation analysis implementation. When considering enrichment score shifts/differences, calculations were done without the log2 transformation.

### Confounders influence analysis

To evaluate the impact of accounting for confounders in the data, differential gene expression was recomputed for all 199 comparisons as described above, without the initial step of searching for confounders. Assuming that all genes found solely when not accounting for confounders are false discoveries, the expected false discovery rate was calculated as the fraction of confounder-associated genes plus a residual 5% of the genes significant in both the differential analysis with and without confounders.

### Counting isoforms in top-down proteomics datasets

We obtained the isoform level supplementary data of Tran *et al*. [40] and Yu *et al*. [41]. For the Yu *et al*. data, we determined the number of isoforms for each protein by counting how many distinct proteoforms were annotated without considering post-translational modifications (PTMs). For the Tran *et al*. data, it is impossible to infer the exact number of isoforms with the information they provide. Instead, we counted the number of genes where isoforms were needed to explain the number of proteoforms annotated (called “species” in this data) given the number of PTMs annotated. Specifically, if the number of annotated proteoforms was larger than the number of possible combinations of PTMs (and no PTMs), we defined the protein as having more than one isoform. Note that the Tran *et al*. result is very conservative and less trustworthy than the Yu *et al*. estimate.

### Enrichment score simulation

We did 1000 simulations of the effect of enrichment of specific types of genes in each of the four following scenarios: 1) enrichment of genes affected by splicing, 2) enrichment of genes affected by expression, 3) enrichment of genes affected by both splicing and expression, 4) genes not enriched for anything (e.i., random). Each simulation had 15,000 genes that could be differentially expressed 10,000 of which could also be differentially spliced (e.i., multi-isoform genes). For each simulation we randomly selected 3000 genes to be differentially expressed and 2000 multi-isoform genes to be differentially spliced. We also randomly sampled a gene-set with 1000 genes to measure enrichment of. To enrich the sampled gene-set for differentially expressed and/or differentially spliced genes we randomly added 1000 genes from the differentially spliced and/or expressed genes. We then calculated the enrichment score for the overlap between the sampled gene-set and the differentially expressed and spliced genes (as described for the pairedGSEA package above). For each of the 4 scenarios we did 1000 simulations where all parameters (except the total number of genes and multi-iso genes) were randomly varied by +/-25% to ensure variability in the simulation. To quantify the difference between the enrichment scores of splicing and expression we subtracted the splicing enrichment score from the gene expression enrichment score.

## Declarations

### Ethics approval and consent to participate

Not applicable

### Consent for publication

Not applicable

### Availability of Data and Materials

All data is available either in the supplementary data or at Zenodo. For each of the 199 comparisons, two RDS objects with the result of running pairedGSEA, using respectively DESeq2/DEXSeq and limma, has been uploaded to Zenodo (DESeq2/DEXSeq based analysis: https://doi.org/10.5281/zenodo.7032090, limma based analysis: https://doi.org/10.5281/zenodo.7866420). These R data object also contains metadata for each sample.

The pairedGSEA R package is available via Bioconductor (https://bioconductor.org/packages/pairedGSEA).

## Author contributions

KVS conceived the study. KVS did the literature survey on the previous use of RNA-seq. SHD and KVS curated the metadata of studies. SHD did all analyses. SHD, LRO, and KVS interpreted the results and wrote the manuscripts.

## Supplementary Figures

**Figure S1:**
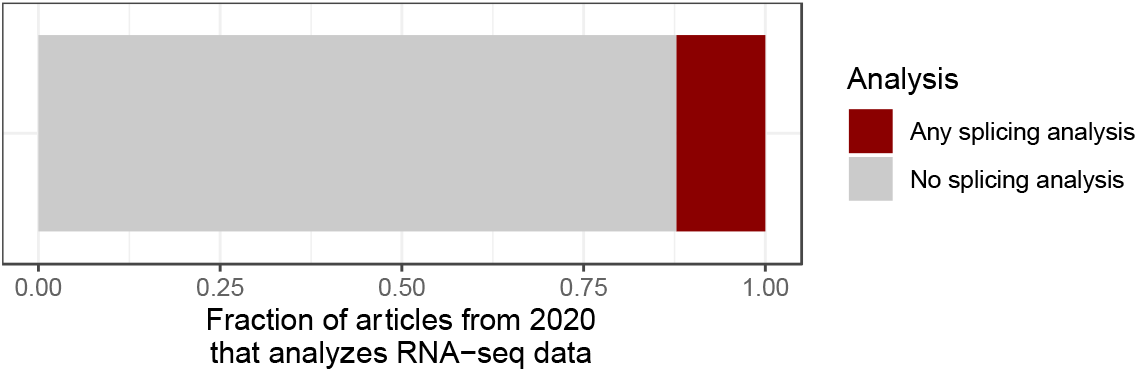
Summary of the manual literature review of articles from 2020 that analyzes RNA-seq data (see Methods).

**Figure S2:**
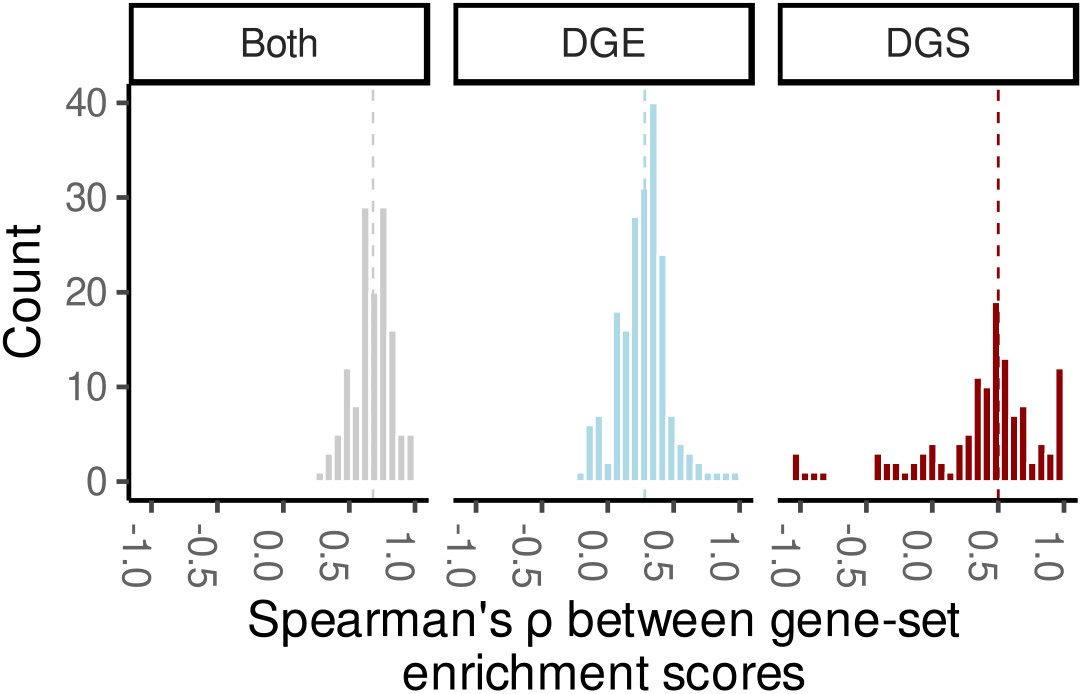
Histogram of the Spearman’s correlations between gene-set enrichment scores for gene sets significantly (FDR-adjusted P-value of <0.05) enriched among differentially expressed or differentially spliced genes. Correlations were computed separately for gene sets enriched among either differentially spliced or expressed genes or both (sub-plots).

**Figure S3:**
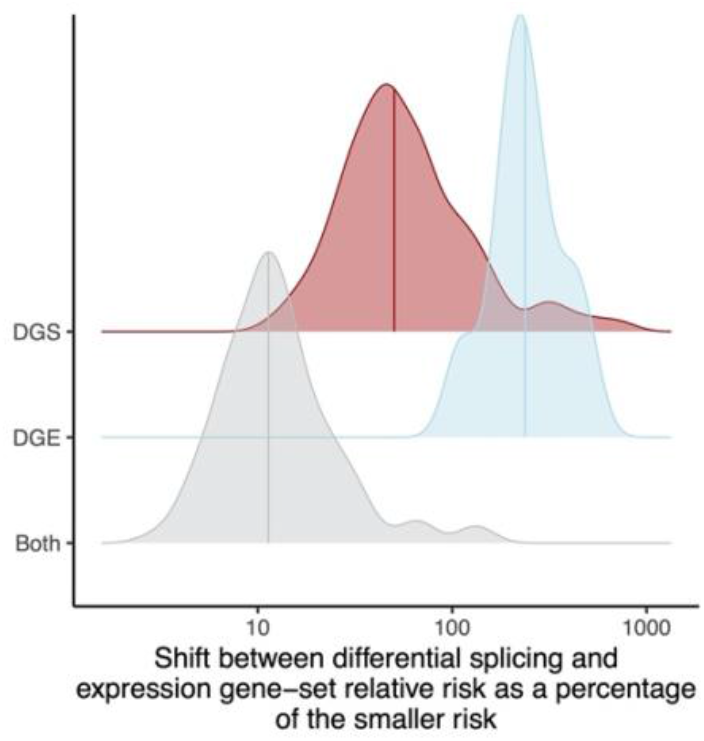
For each comparison and data subset (y-axis), For each comparison, the median differences between relative risks of gene sets enriched among differentially expressed and spliced genes as the percent change of the smallest risk. Data subsets are gene sets that were significantly (FDR-adjusted P-value of <0.05) enriched among differentially spliced genes, expressed genes, or both.

**Figure S4:**
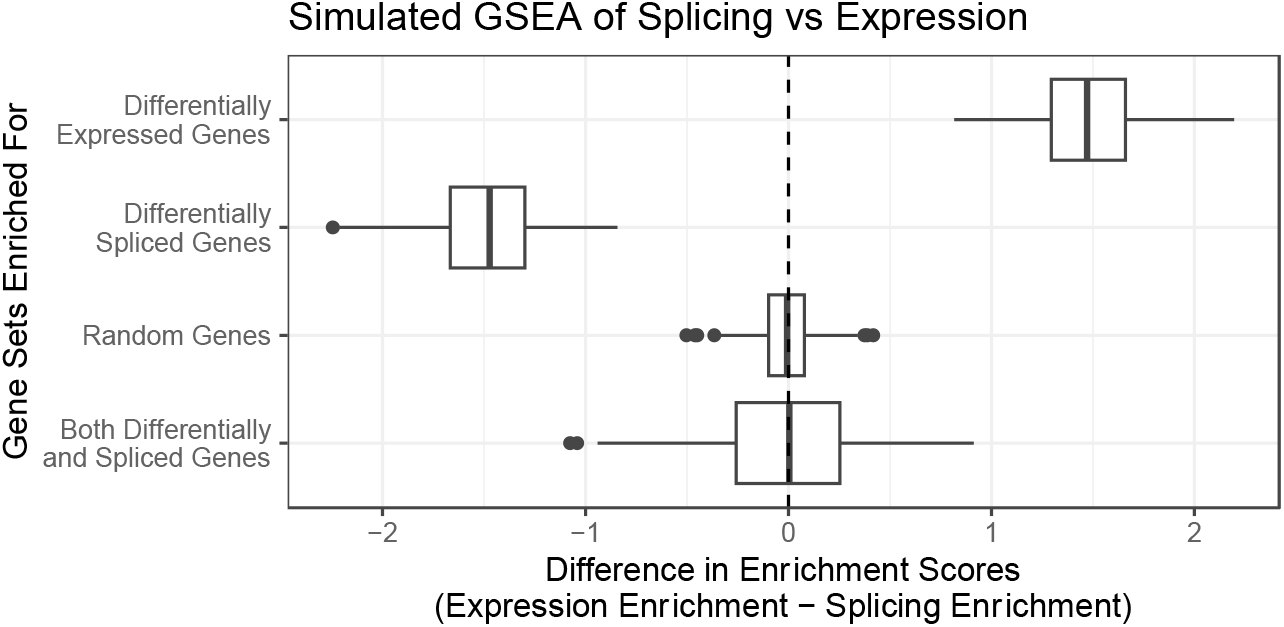
Simulation of how the enrichment score calculated based on gene-set enrichment analysis of both splicing and expression differs (x-axis) depending on which group of genes is enriched (y-axis). Boxplot summarizes 1000 simulations per group.

**Figure S5:**
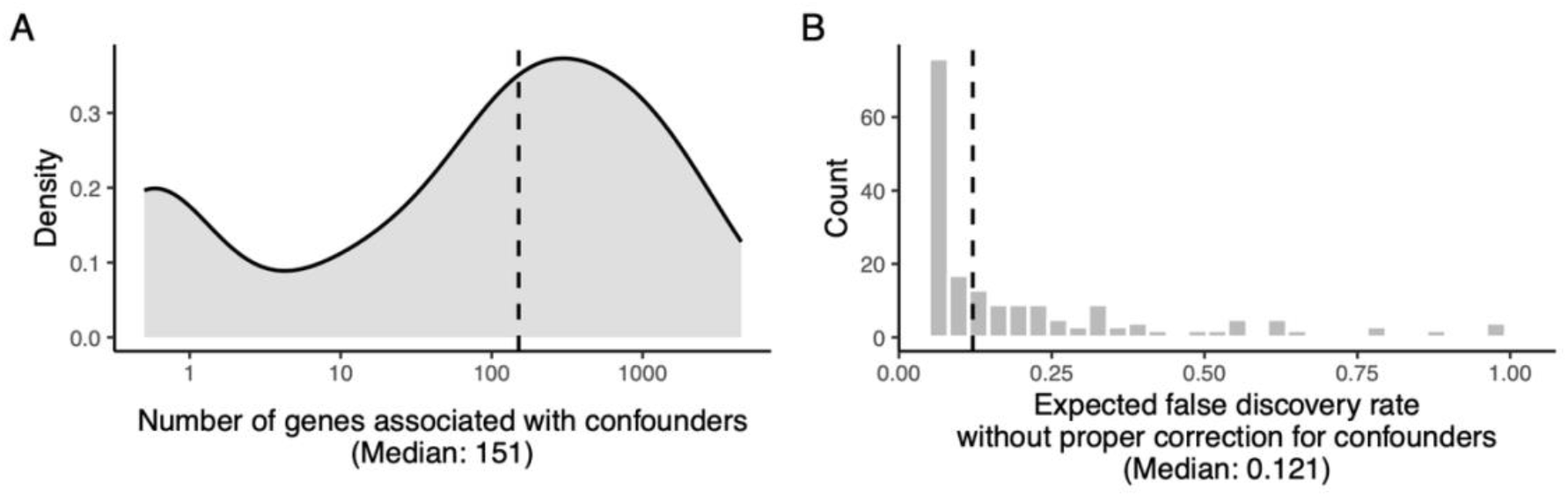
Same as main figure 1B-C but created from the limma based analysis. **A**) The distribution of false positives, i.e., the number of significantly differentially expressed genes only found when not corrected for confounders, across the 199 comparisons. **B**) Histogram of the false discovery rate when not correcting for confounders. Significance is defined as having an FDR-adjusted p-value of <0.05

**Figure S6:**
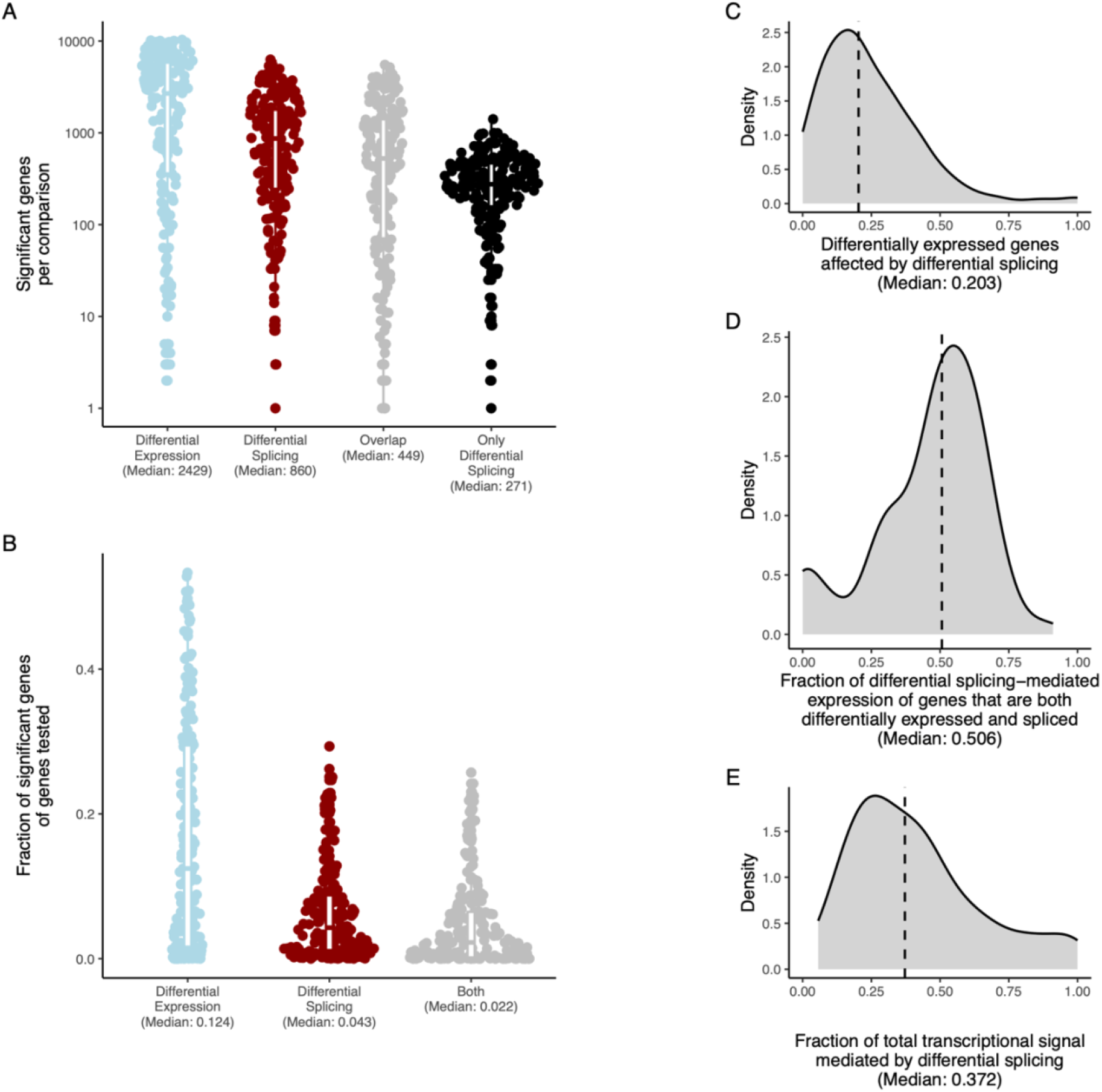
Same as main figure 2 but created from the limma based analysis. Differential Splicing is just as Frequent as Differential Expression. **A**) The number of significant genes for each comparison across analyses.===**B**) For each analysis, the fraction of genes tested that were deemed significant. **C**) The fraction of differentially expressed genes that were also differentially spliced. **D)** Within the genes that are both differentially expressed and spliced, we calculated the fraction of the gene expression that is contributed by differentially spliced transcripts. For each analysis, we extracted the median. **E**) The number of differentially spliced genes as a fraction of the total number of genes either differentially spliced or expressed genes (total transcriptional signal). Across all panes, significance is defined as having an FDR-adjusted P-value of <0.05. Medians are indicated for all plots

